# sPLINK: A Federated, Privacy-Preserving Tool as a Robust Alternative to Meta-Analysis in Genome-Wide Association Studies

**DOI:** 10.1101/2020.06.05.136382

**Authors:** Reza Nasirigerdeh, Reihaneh Torkzadehmahani, Julian Matschinske, Tobias Frisch, Markus List, Julian Späth, Stefan Weiß, Uwe Völker, Dominik Heider, Nina Kerstin Wenke, Tim Kacprowski, Jan Baumbach

## Abstract

Genome-wide association studies (GWAS) have been widely used to unravel connections between genetic variants and diseases. Larger sample sizes in GWAS can lead to discovering more associations and more accurate genetic predictors. However, sharing and combining distributed genomic data to increase the sample size is often challenging or even impossible due to privacy concerns and privacy protection laws such as the GDPR. While meta-analysis has been established as an effective approach to combine summary statistics of several GWAS, its accuracy can be attenuated in the presence of cross-study heterogeneity. Here, we present *sPLINK* (*safe PLINK*), a user-friendly tool, which performs federated GWAS on distributed datasets while preserving the privacy of data and the accuracy of the results. *sPLINK* neither exchanges raw data nor does it rely on summary statistics. Instead, it performs model training in a federated manner, communicating only model parameters between cohorts and a central server. We verify that the federated results from *sPLINK* are the same as those from aggregated analyses conducted with *PLINK*. We demonstrate that *sPLINK* is robust against heterogeneous data (phenotype and confounding factors) distributions across cohorts while existing meta-analysis tools considerably lose accuracy in such scenarios. We also show that *sPLINK* achieves practical runtime, in order of minutes or hours, and acceptable network bandwidth consumption for chi-square and linear/logistic regression tests. Federated analysis with *sPLINK*, thus, has the potential to replace meta-analysis as the gold standard for collaborative GWAS. The user-friendly, readily usable *sPLINK* tool is available at https://exbio.wzw.tum.de/splink.

## 1 Introduction

Genome-wide association studies (GWAS) test millions of single nucleotide polymorphisms (SNPs) to identify possible associations between a specific SNP and disease^1^. They have led to considerable achievements over the past decade including better comprehension of the genetic structure of complex diseases and the discovery of SNPs playing a role in particular traits or disorders^2, 3^. GWAS sample size is an important factor in detecting associations and larger sample sizes lead to identifying more associations and more accurate genetic predictors^2, 4^.

*PLINK*^5^ is a widely used open source software tool for GWAS. The major limitation of *PLINK* is that it can only perform association tests on local data. If multiple cohorts want to conduct a collaborative GWAS to take advantage of larger sample size, they can pool their data for a joint analysis (Figure 1); however, this is close to impossible due to privacy restrictions and data protection issues, especially concerning genetic and medical data. Hence, the field has established methods for meta-analysis of individual studies, where only the results and summary statistics of the individual analyses have to be exchanged^6^ (Figure 1).

**Figure 1.**
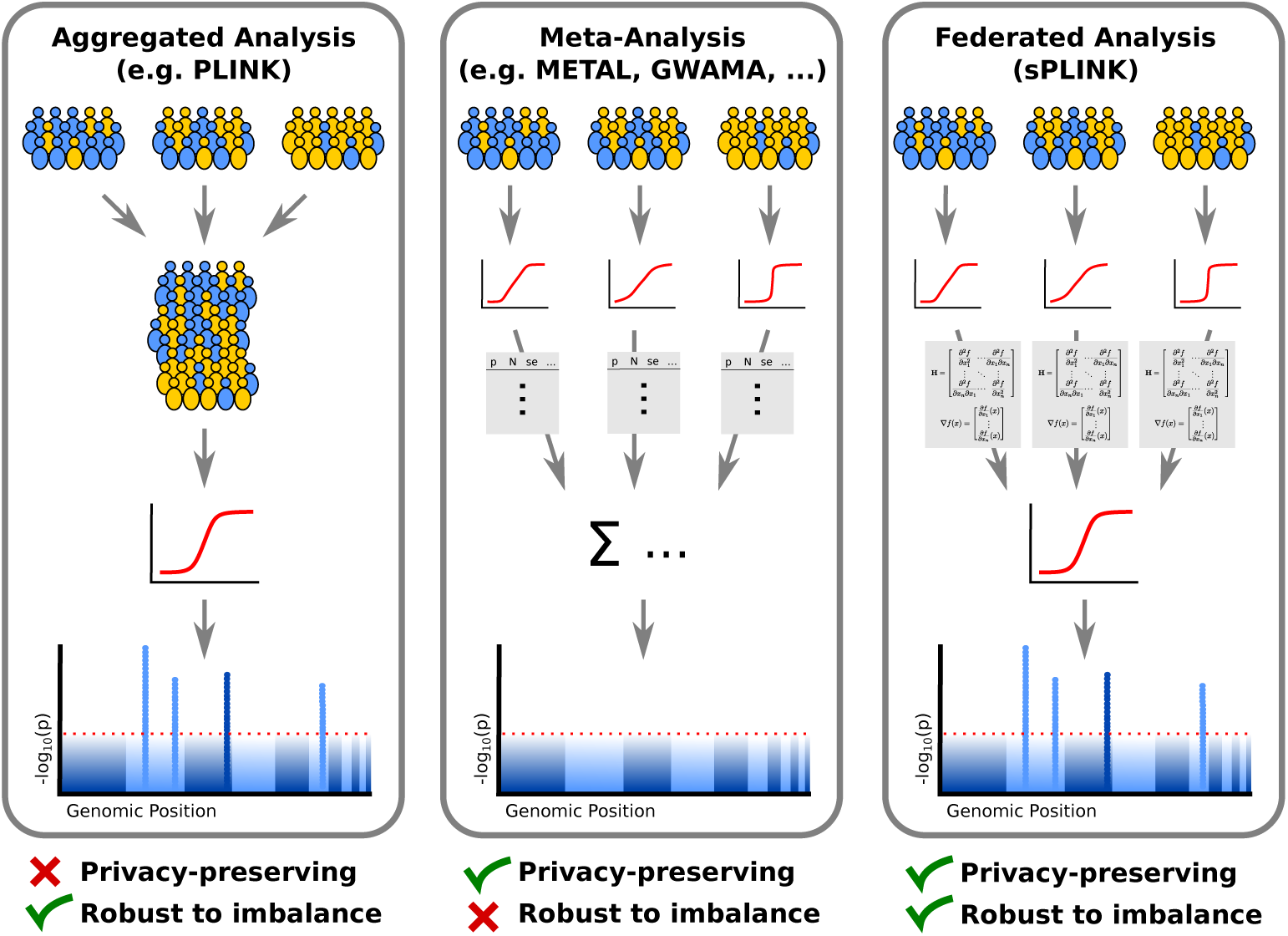
*sPLINK* vs. aggregated analysis and meta-analysis approaches: Aggregated analysis requires cohorts to pool their private data for a joint analysis. Meta-analysis approaches aggregate the summary statistics from the cohorts to estimate the combined p-values. In *sPLINK*, the cohorts extract the model parameters (e.g. Hessian matrices) from the local data, which are aggregated to build a final model. *sPLINK* combines the advantages of aggregated analysis and meta-analysis, i.e. robustness against heterogeneous data and preserving the privacy of cohorts’ data. Yellow/blue color indicates case/control samples.

There are several software packages such as *METAL*^7^, *GWAMA*^8^ and *PLINK*^5^ that implement different meta-analysis models including fixed or random effect models^9^. Although meta-analysis approaches can protect the privacy, they suffer from two main constraints: first of all, they rely on detailed planning and agreement of cohorts on various study parameters such as meta-analysis model (e.g. fixed effect or random effect), meta-analysis tool (e.g. METAL or GWAMA), heterogeneity metric (e.g. Cochran’s *Q* or the *I*^2^ statistic), the covariates to be considered, etc^4^. Most importantly, the statistical power of meta-analysis can be adversely affected in the presence of cross-study heterogeneity, leading to inaccurate estimation of the joint results and yielding misleading conclusions^10, 11^.

To address the aforementioned shortcomings, privacy-preserving collaborative GWAS can be developed using homomorphic encryption^12^(HE), secure multi-party computation^13^(SMPC), and federated learning^14, 15^. In HE, the cohorts encrypt their private data and share it with a single server, which performs operations on the encrypted data from the cohorts to compute the association test results. In SMPC, there are a couple of computing parties and the cohorts extract a separate secret share^16^ (anonymized chunk) from the private data and send it to a computing party. The computing parties calculate intermediate results from the secret shares and exchange the intermediate results with each other. Each computing party computes the final results given all intermediate results. In federated learning, the cohorts extract model parameters (e.g. Hessian matrices) from the private data and share the parameters with a central server. The server aggregates the parameters from all cohorts to calculate the association test results.

*Kamm et al.*^17^ and *Cho et al.*^18^ proposed secure GWAS frameworks based on SMPC. The former developed simple association tests including Cochran–Armitage and chi-square (χ^2^) and the latter implemented only Cochran–Armitage test. Shi et al.^19^ presented a secure SMPC-based logistic regression framework for GWAS. These frameworks inherit the limitations of SMPC itself: They follow the paradigm of “move data to computation”, where they put the processing burden on a few computing parties. Consequently, they are computationally expensive^20^ and are not scalable for large-scale GWAS. Moreover, they suffer from the colluding-parties problem^17^ in which, if the parties send the secret shares of the cohorts to each other, the whole private data of the cohorts is exposed.

*Lu et al.*^21^, *Morshed et al.*^22^, *and Kim et al.*^23^ developed secure chi-square test, linear regression, and logistic regression using HE for GWAS, respectively. Similar to SMPC-based methods, they are not computationally efficient since a single server carries out operations over encrypted data, causing considerable overhead^24^. Additionally, HE-based methods introduce accuracy loss in the association test results^22, 23^. This is because HE only supports addition and multiplication, and as a result, non-linear operations in regression tests should be approximated using those two operations.

To address the limitations of HE/SMPC-based methods, the association tests can be implemented in a federated fashion (Figure 1). Federated learning based methods follow the paradigm of “move computation to data”, distributing the heavy computations among the cohorts while performing lightweight aggregation (simple operations such as addition and multiplication of the parameters) at the central server. There are previous GWAS frameworks^25–27^ based on federated learning but they allow for only one association test (i.e. logistic regression) and/or are not a user-friendly tool set like *PLINK* that can be easily deployed for GWAS.

In this paper, we present a novel federated tool set for GWAS called ***sPLINK (safe PLINK)***. Unlike *PLINK, sPLINK* is applicable to distributed data while protecting the privacy. In *sPLINK*, the private data of cohorts never leaves the site; instead, each cohort installs a local client software that processes local data, extracts the model parameters, and shares only those (but not the data) with the central server (Section 2 and Supplementary B). Contrary to existing HE/SMPC-based methods, *sPLINK* is computationally efficient since heavy computations are distributed across the cohorts while simple aggregation is performed at the server. Compared to the current federated GWAS tools, *sPLINK* not only provides a user-friendly and easy-to-use web interface but also supports multiple association tests including logistic regression^28^, linear regression^29^, and chi-square (*χ*^2^)^30^ for GWAS (Supplementary B).

The advantage of *sPLINK* over current meta-analysis approaches is two-fold: usability and robustness against heterogeneity. *sPLINK* is easier to use for collaborative GWAS compared to meta-analysis. In *sPLINK*, a coordinator initiates a collaborative study and invites the cohorts. The only decision the cohorts make is whether or not to join the study. After accepting the invitation, the cohorts just select the dataset they want to employ in the study (Section 2 and Supplementary A). More importantly, *sPLINK* is robust to data (phenotype and confounding factors) heterogeneity. It gives the same results as aggregated analysis even if the phenotype distribution is imbalanced or if confounding factors are distributed heterogeneously across cohorts. In contrast, meta-analysis tools typically lose statistical power in such imbalanced or heterogeneous scenarios (Figure 2 and Section 3).

**Figure 2.**
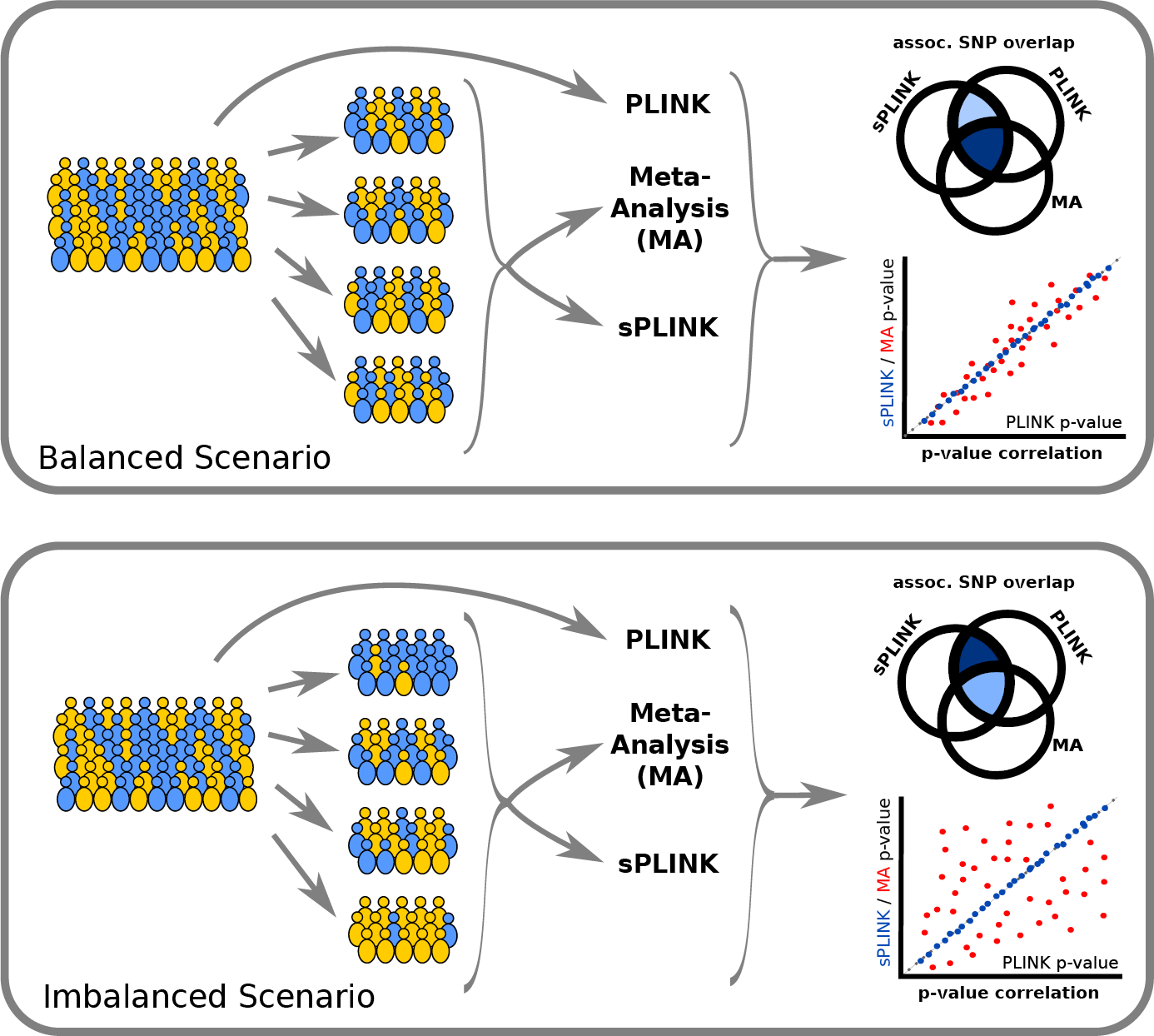
Evaluation of *sPLINK*: We have artificially split datasets into smaller subsets mimicking balanced and imbalanced scenarios. The aggregated analysis using PLINK, meta-analysis with several tools, and results from *sPLINK* have been compared for each scenario. We observe that *sPLINK* agrees with the results of the aggregated analysis in all scenarios, while meta-analyses suffer from imbalanced scenarios.

## 2 Methods

*sPLINK* implements a horizontal (sample-based) federated learning approach^14, 15, 31^ to preserve the privacy of data. Federated learning is a type of distributed learning, where multiple cohorts collaboratively learn a joint (global) model under the orchestration of a central server^32^. The cohorts never share their private data with the server or the other cohorts. Instead, they extract local parameters from their data and send them to the server. The server aggregates the local parameters from all cohorts to compute the global model parameters (or global results), which, in turn, are shared with all cohorts. Specifically, *sPLINK* works with distributed GWAS datasets, where samples are individuals and features are SNPs and categorical or quantitative phenotypic variables. While the samples are different across the cohorts, the feature space is the same since *sPLINK* only considers SNPs and phenotypic variables that are common among all datasets.

The functional workflow of sPLINK is as follows: the coordinator initializes the project, creates a project token for each cohort, and sends the corresponding token to the cohort. Next, the cohorts join the project using their username, password, and token, contributing their dataset to the study. After the cohorts joined, the project is started automatically and the association test results are computed in a federated fashion. When the federated analysis is completed, the cohorts and coordinator can have access to the results (more details on the functional workflow of *sPLINK* can be found in Supplementary A).

From a technical point of view, *sPLINK*’s architecture consists of three main software components: client, server, and web application (WebApp). The client package is installed on the local machine of each cohort with access to the private data. It computes the model parameters from the local data and sends the parameters to the server (Supplementary B). The server and WebApp packages are installed on a central server. The former is responsible for aggregating the local parameters from all the cohorts to calculate the global parameters (Supplementary B). The latter is employed to configure the parameters (e.g. association test) of the new study (Supplementary A). *sPLINK* also provides a chunking capability to handle large datasets containing millions of SNPs. The chunk size (configured by the coordinator) specifies how many SNPs should be processed in parallel. Larger chunk sizes allow for more parallelism, and therefore less running time but require more computational resources (e.g. CPU and main memory) from the local machines of the cohorts and the server. While we provide a readily usable web service running at *exbio* server^1^, the server and WebApp packages can, of course, also be installed by a cohort on an own web server.

## 3 Results

We first verify *sPLINK* by comparing its results with those from aggregated analysis conducted with *PLINK* for all three association tests on a real GWAS dataset from the SHIP study^33^. We refer to this dataset as the *SHIP* dataset, which comprises the records of 3699 individuals with *serum lipase activity* as phenotype. The quantitative version represents the square root transformed serum lipase activity, while the dichotomous (binary) version indicates whether the serum lipase activity of an individual is above or below the upper 25th percentile. The *SHIP* dataset contains around 5 million SNPs and sex, age, smoking status (current-, ex-, or non-smoker), and daily alcohol consumption (in g/day) as confounding factors (Table 1).

**Table 1.**
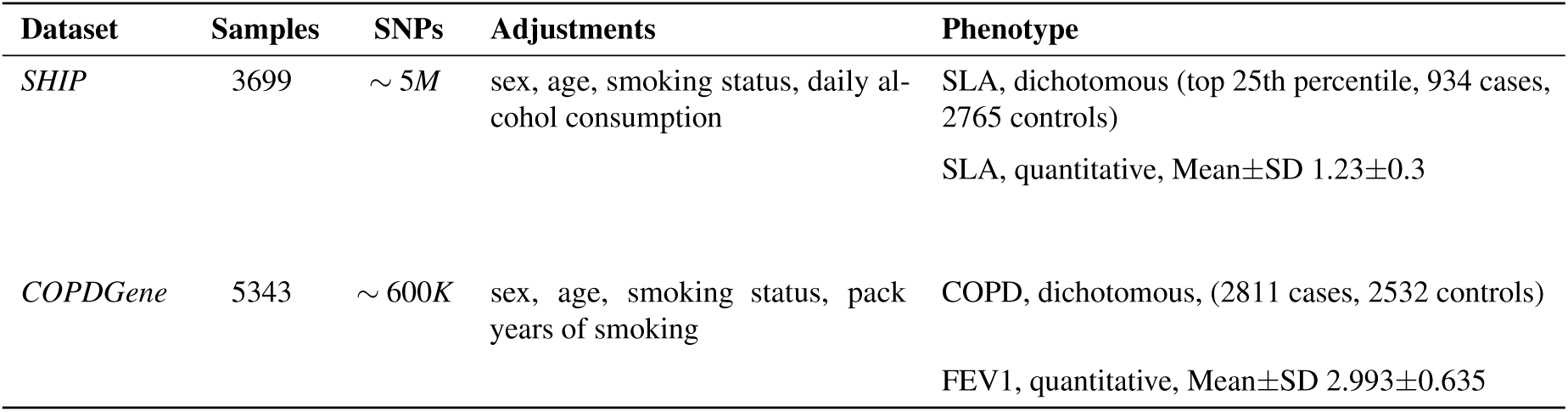
Description of datasets: The *SHIP* (Study of Health in Pomerania) and *COPDGene* (Genetic Epidemiology of COPD) datasets are used to verify *sPLINK* and compare *sPLINK* with the existing meta-analysis tools. COPD: chronic obstructive pulmonary disease, SLA: serum lipase activity, FEV1: forced expiratory volume in one second

We employ the binary phenotype for logistic regression and the chi-square test, and the quantitative phenotype for linear regression. We incorporate all four confounding factors in the regression models and no confounding factor in the chi-square test. We horizontally (sample-wise) split the dataset into four parts, simulating four different cohorts (with sample sizes of 1044, 1006, 941, and 708, respectively). *PLINK* computes the statistics for each association test using the whole dataset while *sPLINK* does it in a federated manner using the splits of the individual cohorts. To be consistent with *PLINK, sPLINK* calculates the same statistics as *PLINK* for the association tests (Supplementary A).

We compute the difference between the p-values (*p*) as well as Pearson correlation coefficient (*ρ*) of p-values from *sPLINK* and *PLINK*. We use -*log*_10_(*p*) since the p-values are typically small and -*log*_10_(*p*) can be a better indicator of small p-value differences. According to Figure 3a-c, the p-value difference is zero for most of the SNPs. We also observe that the maximum difference is 0.162 for a SNP in the linear regression. *sPLINK* and *PLINK* report 4.441 × 10^−16^ and 3.058 × 10^−16^ as p-values for the SNP, respectively. This negligible difference can be attributed to inconsistencies in floating point precision.

**Figure 3.**
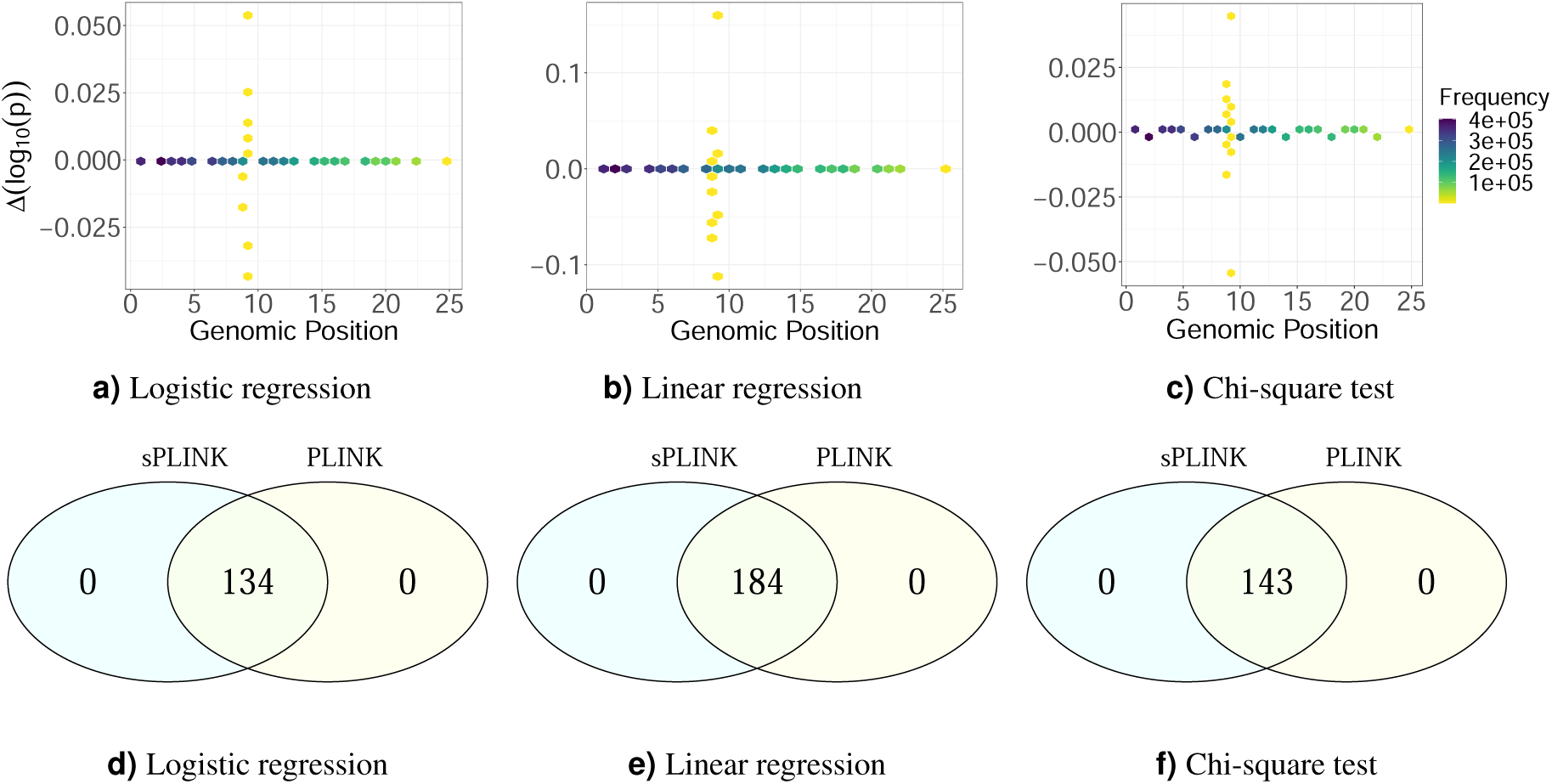
Δ*log*_10_(*p*) between *sPLINK* and *PLINK* as well as the set of SNPs identified by *sPLINK* and *PLINK* as significant for logistic regression (a) and (d), linear regression (b) and (e), and chi-square test (c) and (f), respectively. For most of the SNPs, the difference is zero, indicating that *sPLINK* gives the same p-values as *PLINK*. The negligible difference between p-values for the other SNPs can be attributed to differences in floating point precision. *sPLINK* and *PLINK* also recognize the same set of SNPs as significant.

The correlation coefficient of p-values from *sPLINK* and *PLINK* for all three tests is 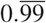, which is consistent with the results of p-value difference from Figure 3a-c. We also investigated the overlap of significantly associated SNPs between *sPLINK* and *PLINK*. We consider a SNP as significant if its p-value is less than 5 × 10^−8^. *PLINK* and *sPLINK* recognize the same set of SNPs as significant (Figure 3d-f). Notably, the identified SNPs have also been implicated in a previous analysis of this dataset^34^. These results indicate that p-values computed by *sPLINK* in a federated manner are the same as those calculated by *PLINK* on the aggregated data (ignoring negligible floating point precision error). In other words, the federated computation in *sPLINK* preserves the accuracy of the results of the association tests.

Next, we compare *sPLINK* with some existing meta-analysis tools, namely *PLINK, METAL*, and *GWAMA*. To do so, we leverage the *COPDGene* dataset (non-hispanic white ethnic group)^35^, which has an equal distribution of case and control samples unlike the *SHIP* dataset (Table 1). *COPDGene* contains 5343 samples (ignoring 1327 samples with missing phenotype value) and around 600K SNPs. We utilized COPD (chronic obstructive pulmonary disease) as the binary phenotype and included sex, age, smoking status, and pack years of smoking as confounding factors^36^ (Table 1).

To simulate cross-study heterogeneity^37^, we consider six different scenarios: *Scenario I* (*Balanced*), *Scenario II* (*Slightly Imbalanced*), *Scenario III* (*Moderately Imbalanced*), *Scenario IV* (*Highly Imbalanced*), *Scenario V* (*Severely Imbalanced*), and *Scenario VI* (*Heterogeneous Confounding Factor*) (Figure 4 and Figure 5). In each scenario, we partition the dataset into three splits with the same sample size of 1781. The distribution of all four confounding factors is homogeneous (similar) across the splits for the first five scenarios. The splits have the same (and balanced) case-control ratio in *Scenario I* and *Scenario VI* but their case-control ratio is different for the imbalanced scenarios (Figure 4). In *Scenario VI*, the values of two confounding factors (i.e. smoking status and age) are homogeneously distributed among the splits; however, the distribution of sex and pack years of smoking is slightly and highly heterogeneous across the splits, respectively (Figure 5).

**Figure 4.**
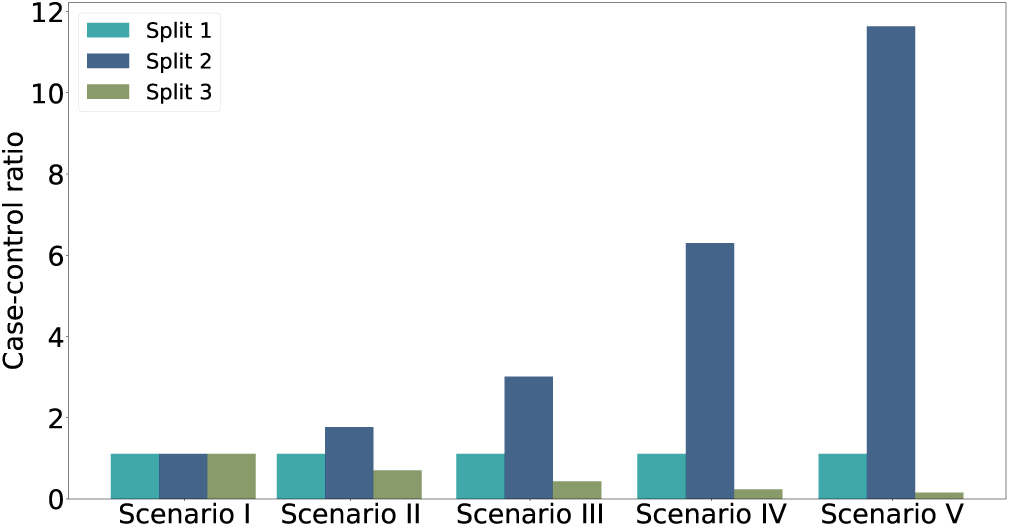
Scenario I-V: The case-control ratio is the same (1.11) for all splits in the balanced scenario (I) while the splits have different case-control ratio in the imbalanced scenarios (II-V). The distribution of all four confounding factors is homogeneous across the splits.

**Figure 5.**
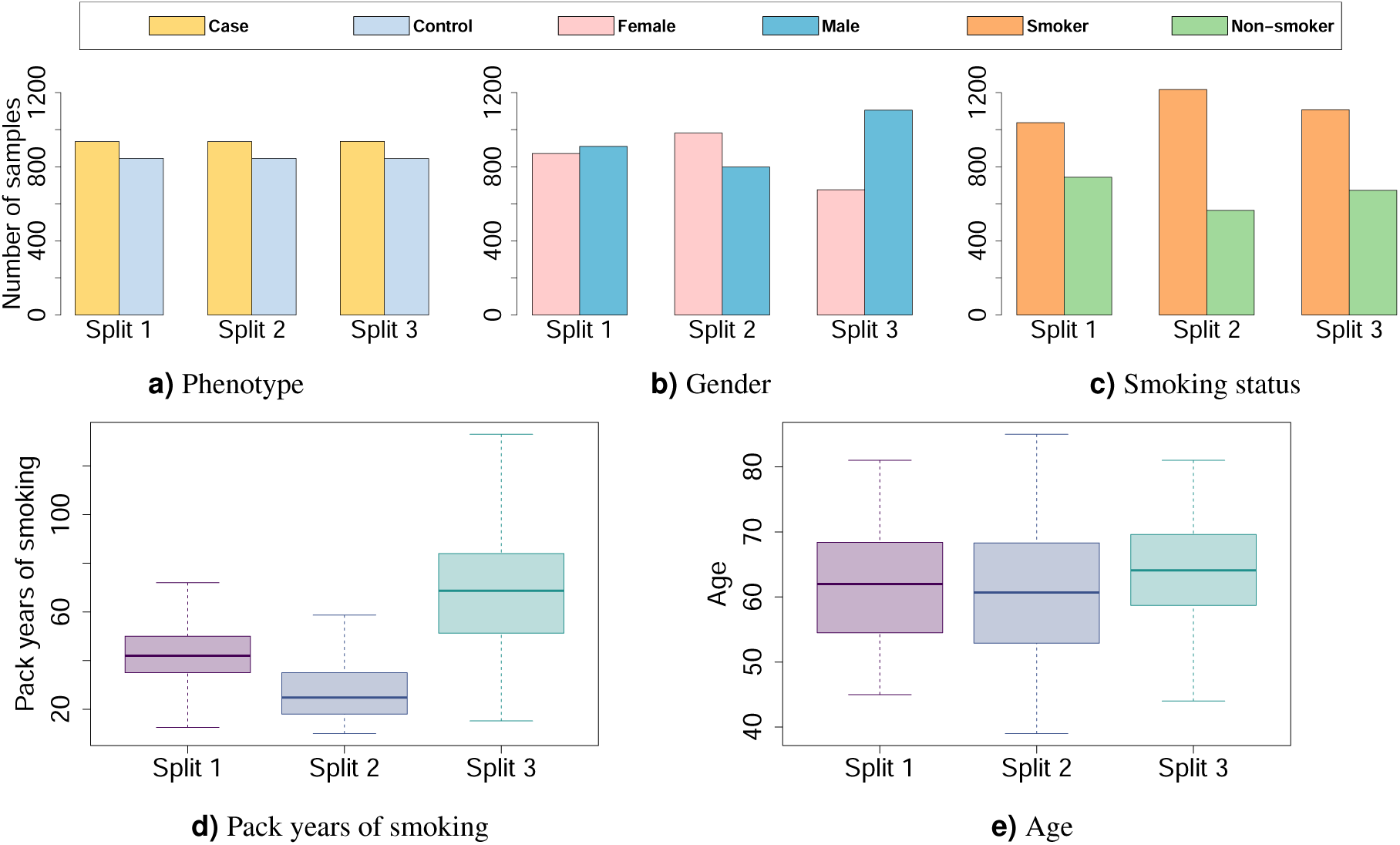
Scenario VI (Heterogeneous Confounding Factor): The phenotype distribution is the same and balanced; the values of smoking status and age are homogeneously distributed; the distribution of gender and pack years of smoking are slightly and highly heterogeneous across the splits, respectively.

We obtain the summary statistics (e.g. minor allele, odds ratio, standard error, etc.) for each split to conduct meta-analyses. The results are then compared to the federated analysis employing *sPLINK*. Figure 6a shows the Pearson correlation coefficient of -*log*_10_(*p*) between each tool and the aggregated analysis for all six scenarios. Figure 6b depicts the number of SNPs correctly identified as significant by the tools (true positives). We repeat each scenario five times, where in each run the dataset is randomly partitioned into three splits (the same sample size) according to the case-control ratio and confounding factor distributions as depicted in Figure 4 and Figure 5.

**Figure 6.**
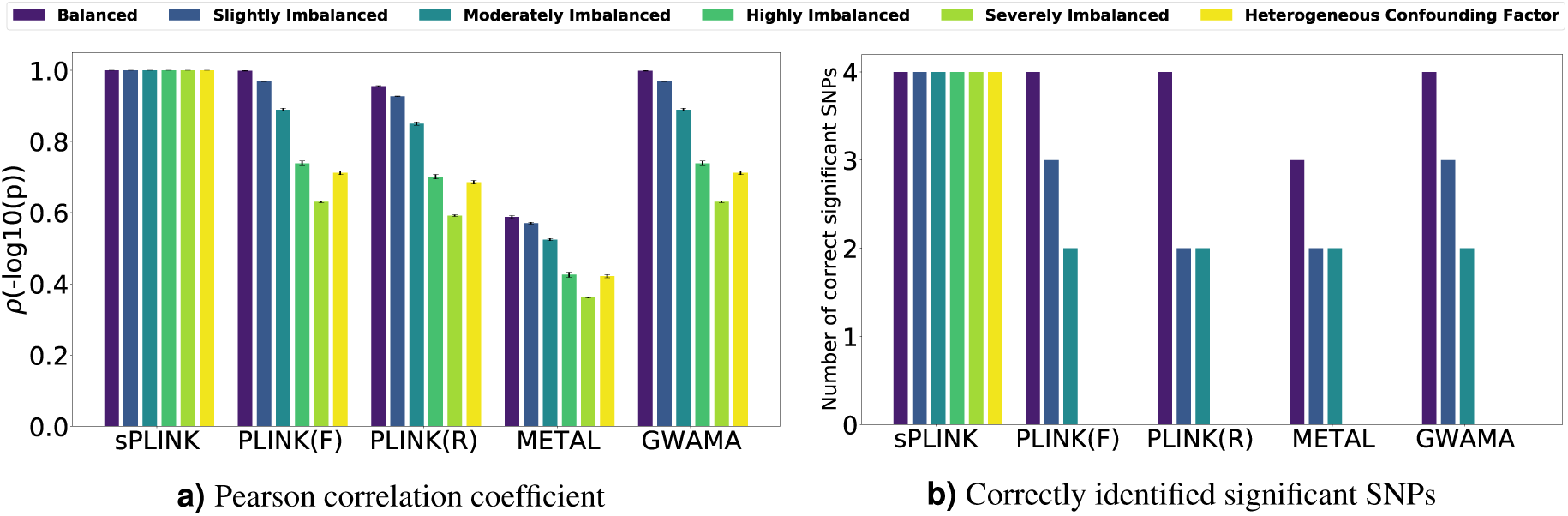
(a) Pearson correlation coefficient (*ρ*) of -*log*_10_(*p*) between each tool and aggregated analysis and (b) the number of SNPs correctly identified as significant (true positives) by each tool. The height of each bar in (a) and (b) represents the average and median across the five runs, respectively. Error bars indicate the standard deviation. *F* and *R* stand for fixed-effect and random-effect, respectively.

According to Figure 6a, the correlation of p-values between *sPLINK* and the aggregated analysis is ∼1.0 for all six scenarios, implying that *sPLINK* gives the same p-values as aggregated analysis regardless of how phenotypes or confounding factors have been distributed across the cohorts. In contrast, the correlation coefficient for the meta-analysis tools shrinks with increasing imbalance/heterogeneity, indicating loss of accuracy. Figure 6b illustrates that *sPLINK* correctly identifies all four significant SNPs in all scenarios. In the balanced scenario, almost all meta-analysis tools perform well and recognize all significant SNPs. An exception is *METAL*, which misses one of them. However, they miss more and more significant SNPs as the phenotype imbalance across the splits increases. In the *Highly Imbalanced* and *Severely Imbalanced* scenarios, the meta-analysis tools cannot recognize any significant SNP. This is also the case if the distribution of some confounding factors becomes heterogeneous across the cohorts (*Scenario VI*). We checked the number of SNPs wrongly identified as significant by the tools (false positives) too. *sPLINK* has no false positive in any of the scenarios and the meta-analysis tools introduce zero or one false positive depending on the scenario.

Finally, we measure the runtime and network bandwidth consumption of *sPLINK* (Supplementary C) using COPDGene dataset (5343 samples and ∼ 600*K* SNPs) partitioned into three splits of the same sample size. *sPLINK* calculates the test results in 22 *min*, 56 *min*, and 3.5 *h*, exchanging total of 0.46 GB, 0.9 GB, and 8.33 GB traffic for chi-square, linear regression, and logistic regression, respectively. This indicates that *sPLINK* achieves practical running time and acceptable network bandwidth usage for all three association tests. More details and comparison between *sPLINK* and SMPC-based approaches can be found in Supplementary C.

Table 2 shows a concise comparison between *sPLINK* and state-of-the-art approaches. Unlike *PLINK, sPLINK* protects the privacy of the cohorts’ data. *sPLINK* is also robust against the imbalance/heterogeneity of phenotype/confounding factor distributions across the cohorts. That is, *sPLINK* always delivers the same p-values as aggregated analysis and correctly identifies all significant SNPs independent of the phenotype or confounding factor distribution in the cohorts. In contrast, the meta-analysis tools lose their statistical power in imbalanced phenotype scenarios, leading to missing some or all significant SNPs. This is also the case even if the phenotype distribution is balanced but the values of confounding factor(s) have heterogeneously been distributed across the datasets. Compared to the existing SMPC, HE, or federated learning based approaches, *sPLINK* is computationally efficient and supports multiple association tests including chi-square and linear/logistic regression. Note that we explored the robustness to heterogeneity systematically only for tools offering chi-square, linear and logistic regression models.

**Table 2.** Comparison between *sPLINK* and state-of-the-art approaches.

| Tool/Study | Privacy | Robustness to heterogeneity | Computationally efficient | Linear regression | Logistic regression |
| --- | --- | --- | --- | --- | --- |
| *PLINK* | ✗ | ✓ | ✓ | ✓ | ✓ |
| *Meta-analysis* | ✓ | ✗ | ✓ | ✓ | ✓ |
| *Kamm et al.*<sup>17</sup> | ✓ | - | ✗ | ✗ | ✗ |
| *Cho et al.*<sup>18</sup> | ✓ | - | ✗ | ✗ | ✗ |
| *Morshed et al.*<sup>22</sup> | ✓ | - | ✗ | ✓ | ✗ |
| *Kim et al.*<sup>23</sup> | ✓ | - | ✗ | ✗ | ✓ |
| *GLORE*<sup>25</sup> | ✓ | - | ✓ | ✗ | ✓ |
| *sPLINK* | ✓ | ✓ | ✓ | ✓ | ✓ |

## 4 Discussion and Conclusion

We introduced *sPLINK*, a user-friendly federated tool set for GWAS. *sPLINK* preserves the privacy of the cohorts’ data without sacrificing the accuracy of the test results. It supports multiple association tests including chi-square, linear regression, and logistic regression. *sPLINK* is consistent with *PLINK* in terms of the input data formats it supports and the results it reports to the user. We compared *sPLINK* to aggregated analysis with *PLINK* as well as meta-analysis with *METAL, GWAMA*, and *PLINK*. While *sPLINK* is robust against the heterogeneity of phenotype or confounding factor distributions across separate datasets, the statistical power of the meta-analysis tools is attenuated in imbalanced/heterogeneous scenarios. We argued that *sPLINK* is easier to use for collaborative GWAS compared to meta-analysis approaches thanks to its straightforward functional workflow. We also showed that *sPLINK* achieves practical runtime, in order of minutes or hours, and acceptable network usage for the association tests it supports.

The future development of *sPLINK* can go in several directions. We plan to implement the federated version of more association tests^38^ or more machine learning algorithms including random forest^39^ or deep neural networks (DNN)^40^ leveraged by the GWAS community in *sPLINK*. We will also investigate *sPLINK*’s potential to tackle other open challenges in GWAS such as trans-ethnicity^41^, where the samples in the distributed datasets are from different ethnic groups.

In conclusion, *sPLINK* is a novel and robust alternative to meta-analysis, which performs collaborative GWAS in a federated and privacy-preserving manner. It has the potential to immensely impact the statistical genetics community by addressing current challenges in GWAS including cross-study heterogeneity and, thus, to replace meta-analysis as the gold standard for collaborative GWAS.

## Acknowledgments

This project has received funding from the European Union’s Horizon 2020 research and innovation programme under grant agreement No 826078. This reflects only the author’s view and the European Commission is not responsible for any use that may be made of the information it contains. This work was supported by the BMBF-funded de.NBI Cloud within the German Network for Bioinformatics Infrastructure (de.NBI) (031A537B, 031A533A, 031A538A, 031A533B, 031A535A, 031A537C, 031A534A, 031A532B). SHIP is part of the Community Medicine Research net of the University of Greifswald, Germany (www.community-medicine.de), which is funded by the Federal Ministry of Education and Research (grants no. 01ZZ9603, 01ZZ0103, and 01ZZ0403), the Siemens AG, the Ministry of Cultural Affairs as well as the Social Ministry of the Federal State of Mecklenburg-West Pomerania, and the network ‘Greifswald Approach to Individualized Medicine (GANI_MED)’ funded by the Federal Ministry of Education and Research (grant 03IS2061A). ExomeChip data have been supported by the Federal Ministry of Education and Research (grant no. 03Z1CN22) and the Federal State of Mecklenburg-West Pomerania

## A Functional workflow

The *functional workflow* of *sPLINK* (Figure 7) is comprised of the following steps:

**Figure 7.**
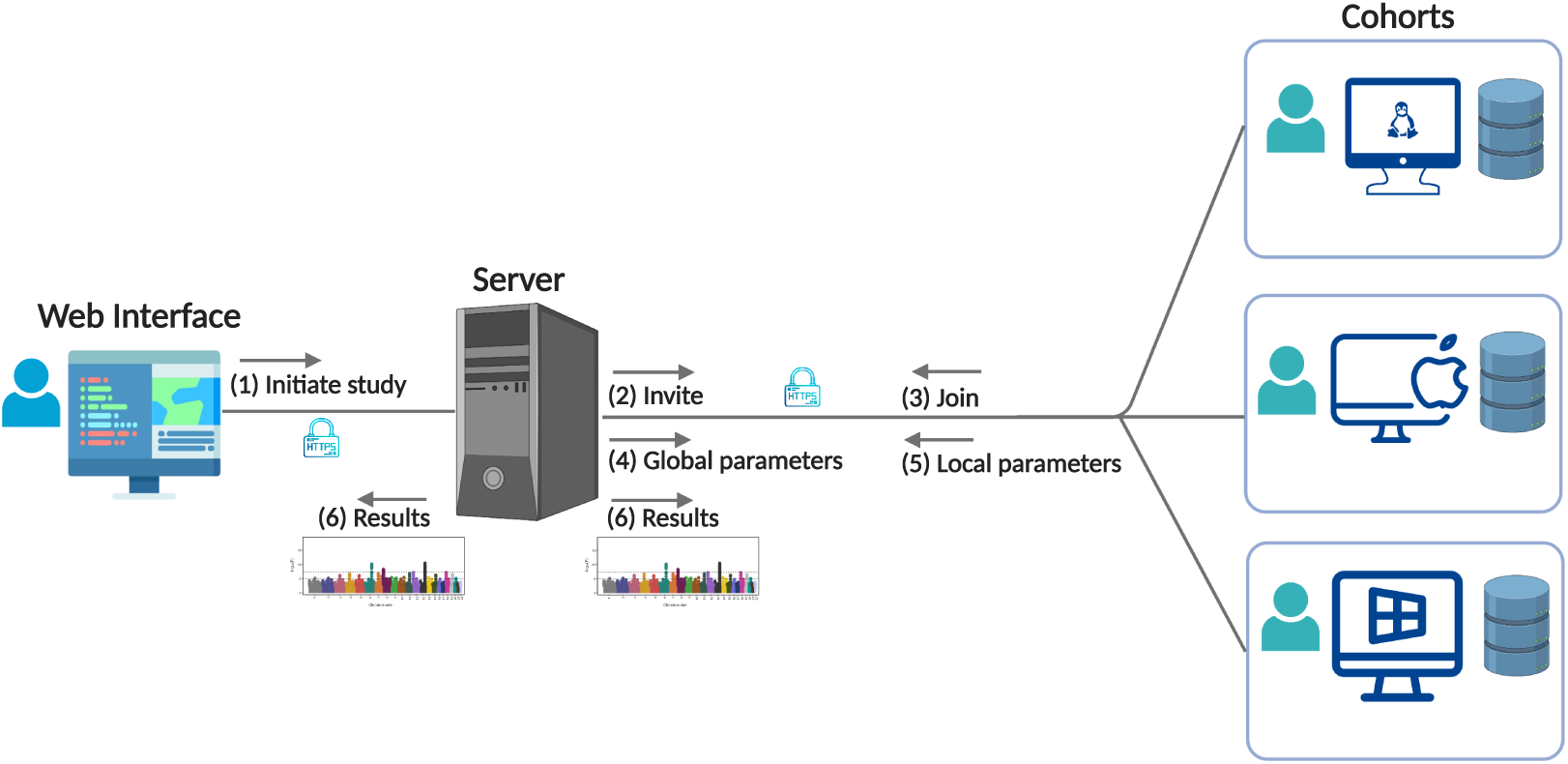
Functional workflow of *sPLINK*: the coordinator creates a new project and invites a set of cohorts to join the project. The cohorts join the project and select the dataset. The project is started automatically, when all cohorts joined. After the computation is done, the cohorts and coordinator can access the results. All communications are performed in a secure channel over HTTPS protocol. The cohorts can use Linux distributions, Microsoft Windows, or MacOS to run the client package.

1. **Project creation**: The coordinator creates the project (new study) through the Web interface. To this end, he/she specifies project name, association test name, chunk size, and the list of confounding features (only for regression tests). He/She also generates a unique project token for each cohort.
2. **Cohort Invitation**: The server sends the project token to each cohort for inviting them to the project.
3. **Cohort joining**: The cohorts use their corresponding username, password, and project token to join the project. Next, they choose the dataset they want to employ in the study. To be consistent with *PLINK, sPLINK* supports .*bed* (value of SNPs), .*fam* (sample IDs as well as gender and phenotype values), .*bim* (chromosome number, name, and base-pair distance of each SNP), .*cov* (value of confounding factors), and .*pheno* (phenotype values that should be used instead of those in .*fam* file) file formats as specified in the *PLINK* manual^42^. For linear regression, phenotype values must be quantitative while for logistic regression and chi-square, phenotype values have to be binary (control/case are encoded as 1/2).
4. **Federated computation**: In *sPLINK*, the association test results are computed by the client package (running on the local machines of cohorts) and server package (running in the central server) in a federated manner. The computation is iterative and consists of four general steps:
  a. **Get global parameters**: All clients obtain the required global model parameter values from the server.
  b. **Compute local parameters**: Each client computes the local model parameter values using the local data and global parameter values from the server.
  c. **Post local parameters**: The local parameter values are shared with the server.
  d. **Aggregate local parameters**: When the server receives the local parameters from all clients, it starts the aggregation process. This involves simple logical operations such as equality check or mathematical operations like addition, subtraction, or multiplication of scalar values or matrices. The results of the aggregation process are the global model parameter values.
5. **Result downloading**: The final results are automatically downloaded for the cohorts but the coordinator needs to download them manually through the web interface. Similar to *PLINK, sPLINK* reports minor allele name (*A1*) and p-value (*P*) for all three association tests, chi-square (*CHISQ*), odds ratio (*OR*), minor allele frequency in cases (*F*_*A*), and minor allele frequency in controls (*F*_*U*) for chi-square test, and the number of non-missing samples (*NMISS*), beta (*BETA*), and t-statistic (*STAT*) for linear and logistic regression tests.

## B Computational workflow

The *computational workflow* of *sPLINK* involves six steps in common among all association tests as well as a couple of steps specific to each association test (Figure 8). In the following, we provide an overview of each step.

**Figure 8.**
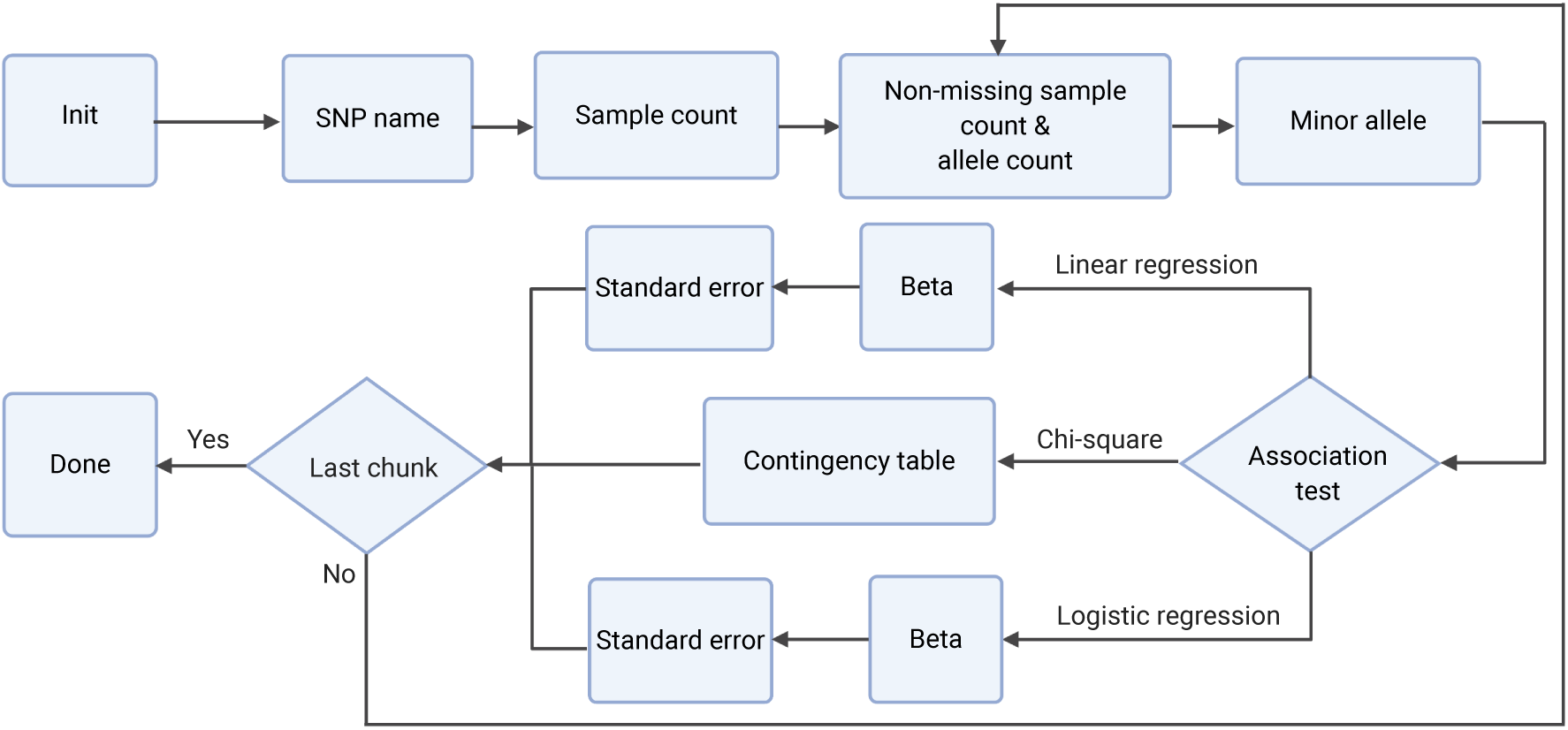
Computational workflow of *sPLINK*: six steps are common among all three association tests: (1) *Init* in which the clients open the required files and prepare the dataset for computation; (2) *SNP name*, where the clients share the SNP names from .bim file with the server; (3) *Sample count* in which each client sends the number of the samples in the dataset considering those with missing values to the server; (4) *Non-missing sample count* & *allele count*, where the clients share the sample count ignoring those with missing values as well as the frequency of each allele with the server; (5) *Minor allele* in which the clients update the minor allele name based on global minor allele and update the mapping of the SNP values accordingly; after the *Minor allele* step, there are association test specific steps: *Beta* and *Standard error* steps for regression tests and *Contingency table* for chi-square test. Logistic and linear regression compute beta and standard error in different ways, e.g. the *Beta* step is iterative for logistic regression. The regression tests compute p-values from standard error values while chi-square does it using the contingency table. The last step is *Done*, where the results from federated computation are shared with the cohorts. Notice that the results from *Non-missing sample count* & *allele count, Minor allele*, and association test specific steps are per SNP and they are computed chunk by chunk.

1. **Init**: Each client *i* opens the files of the dataset selected by the cohort to be employed in the study and creates its phenotype vector (*Y*_*i*_) and feature matrix (*X*_*i*_), which includes the value of SNPs and confounding factors. There is a separate feature matrix for each SNP but the phenotype vector is the same for all SNPs. Assume a dataset containing three SNPs named *SNP1, SNP2*, and *SNP3* and *age* and *sex* as confounding features. There will be three different feature matrices, one feature matrix per SNP. For instance, the feature matrix of *SNP1* has three columns including *SNP1, age*, and *sex* values. Phenotype vector and feature matrix are the private data of the cohorts. They cannot be shared with the coordinator or the other cohorts. The aggregation process in the server just makes sure that all clients successfully initialized the project.
2. **SNP name**: Each client extracts the SNP names from its .*bim* file. In the aggregation process, the server computes the intersection of all SNP names. Only shared SNPs are considered in the computation of the association test results.
3. **Sample count**: Each client *i* calculates its local sample count *n*_*i*_ (number of samples in its dataset including missing samples, which is the size of vector *Y*_*i*_). The server adds up the local sample counts from *K* clients to compute the global sample count: 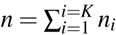
4. **Non-missing sample count** & **allele count**: In this step, SNPs are split into chunks which can be processed in parallel. The chunking capability is provided to handle very large datasets containing millions of SNPs. The clients compute the sample count after filtering out the missing values (value of -9 is considered as missing). Likewise, they calculate the local allele count by counting the number of alleles in each SNP. In the aggregation process, the server adds up the local non-missing sample count from the clients to compute the global non-missing sample count. Next, it does the same for the local allele count and calculates the global allele count. Finally, the server determines the global minor allele based on the value of the global allele count.
5. **Minor allele**: The clients compare their local minor allele with the global minor allele. If they are the same, they do nothing. Otherwise, they update the mapping of SNP values read from .bed file. Each SNP value can be 0,1,2 or 3 (missing value). These values are encoded based on the minor allele name. If the minor allele is changed, the value of the SNP needs to be swapped if it is 0 or 2. Thus, if a client’s minor allele is different from global minor allele, it inverses the mapping of SNP values (0 → 2 and 2 → 0). The aggregation in the server makes sure that all clients successfully completed this step.
6. **Association test specific steps**: In the following, we elaborate on the steps specific to each association test. Regarding regression tests, *sPLINK* implements the federated versions of ordinary least squares (OLS) linear regression and Newton-Raphson method based logistic regression. **Chi-square:** The only test-specific step for chi-square test is *Contingency table*, where each client *i* computes its local contingency table containing minor allele frequency for cases (*p*_*i*_), minor allele frequency for controls (*r*_*i*_), major allele frequency for cases (*q*_*i*_), and major allele frequency for controls (*s*_*i*_). The server adds up the locally computed contingency tables from *K* clients to compute the global (observed) contingency table (Table 3). It also calculates an expected contingency table based on the observed contingency table (Table 4).

**Table 3.** Global (observed) contingency table

|  | Minor allele | Major allele | Total |
| --- | --- | --- | --- |
| Case | $p = \sum_{i=1}^{i=K} p_i$ | $q = \sum_{i=1}^{i=K} q_i$ | $p + q$ |
| Control | $r = \sum_{i=1}^{i=K} r_i$ | $s = \sum_{i=1}^{i=K} s_i$ | $r + s$ |
| Total | $p + r$ | $q + s$ | $n$ |

**Table 4.** Expected contingency table Given the observed contingency table (*O*) and the expected contingency table (*E*), the server computes odds ratio (OR), *χ*^2^, and p-value (P) as follows:

|  | Minor allele | Major allele |
| --- | --- | --- |
| Case | $\frac{(p+q) \times (p+r)}{n}$ | $\frac{(p+q) \times (q+s)}{n}$ |
| Control | $\frac{(r+s) \times (p+r)}{n}$ | $\frac{(r+s) \times (q+s)}{n}$ |

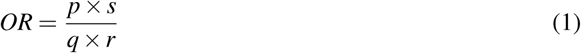

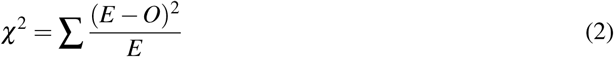

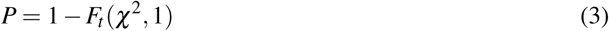

where *F*_*t*_ is the cumulative distribution function (CDF) of *χ*^2^ distribution (degree of freedom is 1). **Linear regression:** *Beta* and *Standard error* are two steps specific to linear regression test. In the *Beta* step, each client *i* computes 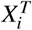 and 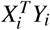, where 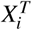 is the transpose of *X*_*i*_. In the aggregation process, the server performs the following calculations (*K* is the number of clients):

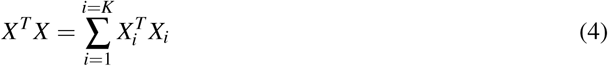

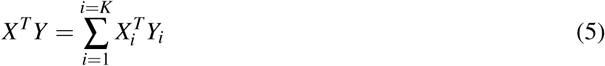

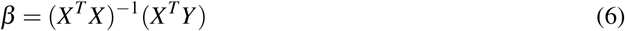

where ()^−1^ indicates the inverse matrix. In the *Standard error* step, each client *i* calculates the local sum square error (*SSE*_*i*_) by having the global *β* vector.

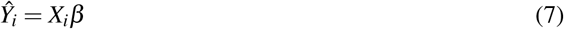

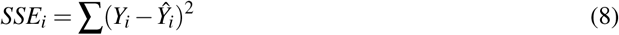

and then the server calculates the global standard error vector (SE) as follows:

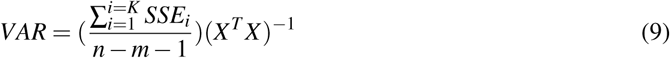

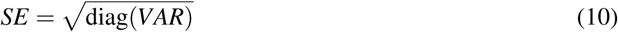

where *n* is global non-missing sample count, *m* is the number of features (1 + number of confounding factors), and *diag* is the main diagonal of the matrix. Given the standard error vector, the server computes the *T statistic* (*T*) and *p-value* (*P*) as follows:

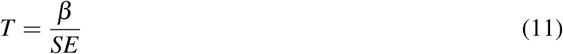

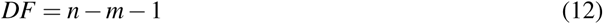

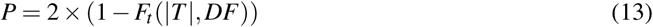

in which *DF* is degree of freedom and *F*_*t*_ is the CDF of T distribution. **Logistic regression:** Similar to linear regression, logistic regression has two specific steps: *Beta* and *Standard error*. However, the *Beta* step is iterative in logistic regression (maximum number of iterations is specified by the coordinator and its default value is 20). In each iteration, each client *i* computes local gradient (∇_*i*_), Hessian matrix (*H*_*i*_) and log-likelihood (*L*_*i*_) as follows:

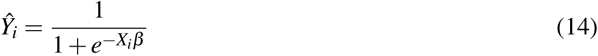

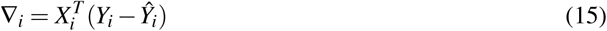

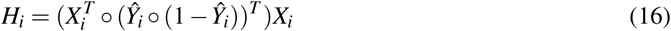

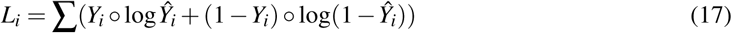

where *β* is the global beta vector from the previous iteration and ∘ indicates element-wise multiplication. The server aggregates the local gradients, Hessian matrices and log-likelihood values from *K* cohorts as follows:

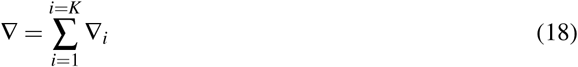

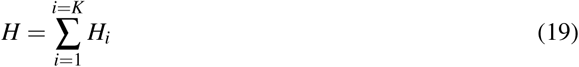

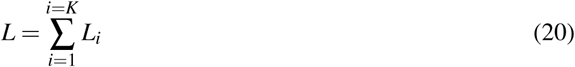

Then, it updates the *β* values accordingly:

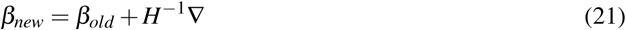

where *β*_*old*_ is the *β* value from the previous iteration. The server also compares the newly computed log-likelihood value (L) with the one from the previous iteration(*L*_*old*_). A difference less than a pre-specified threshold indicates that the *β* values have converged. In the *Standard error* step, the server shares the global *β* values with the clients. Each client *i* computes its local Hessian matrix (*H*_*i*_) using the global *β*. The server gets the local Hessian matrices from *K* cohorts and applies the following formula to obtain the global standard error vector (SE):

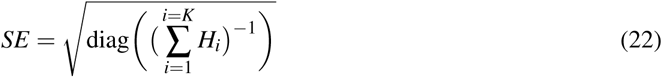

Having standard error values, the server calculates T statistics (*T*) and p-value (*P*) as follows:

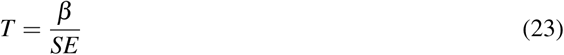

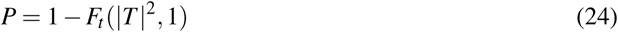

where *F*_*t*_ is CDF of *χ*^2^ distribution (degree of freedom is 1).
7. **Done**: The computation of association test results have been completed for all chunks and the results are shared with all cohorts.

## C Runtime and network bandwidth usage

We measure the runtime (Figure 9a) and network bandwidth usage (Figure 9b) of *sPLINK* for each association test using COPDGene dataset (5343 samples and ∼ 600*K* SNPs) partitioned into three splits of the same sample size (1781). We use *COPD* in chi-square as well as logistic regression and *FEV1* in linear regression as phenotype. We include age, sex, smoking status, and pack years of smoking as confounding factors only for the regression tests. The server and WebApp packages are running in a docker container (8 CPU cores and 32 GB of main memory allocated) at *exbio* server. Three commodity laptops (4 cores and 16 GB of main memory) located at *Munich* or *Freising* are running the client package and host the splits. They communicate with the server through Internet with connection speeds of 100, 50, and 50 Mbps (megabits per second), respectively.

**Figure 9.**
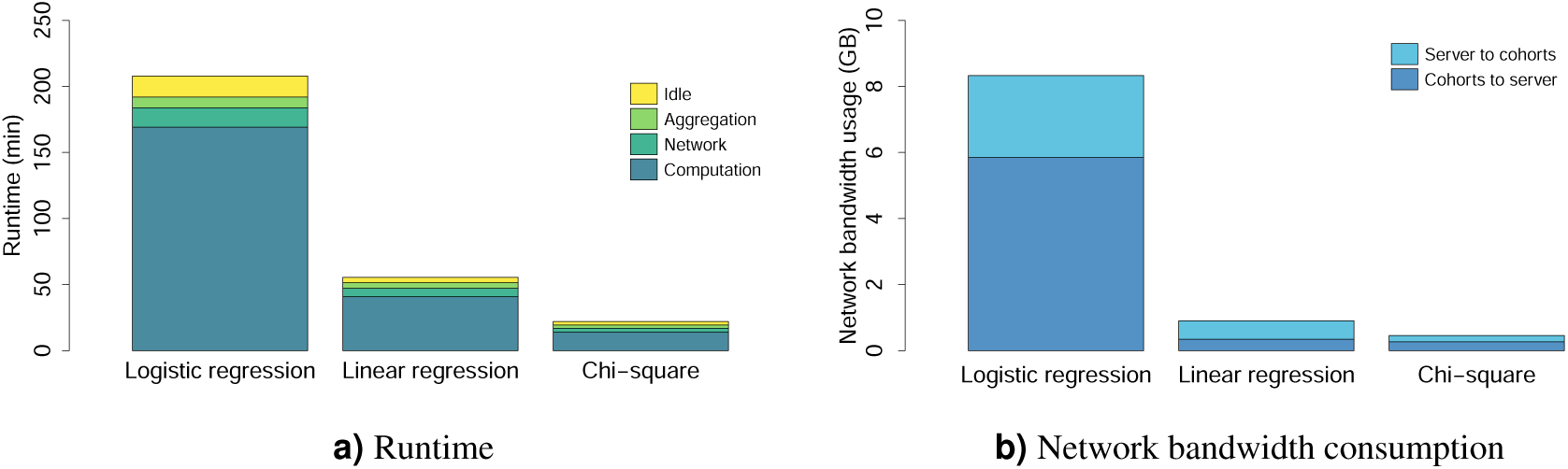
Runtime of *sPLINK* (a) consists of processing time at cohorts, network communication time, aggregation time at the server, and idle time. The computation time contributes the most in the runtime. The network bandwidth consumption of *sPLINK* (b) is mainly due to sending model parameter values from the cohorts to the server.

Figure 9a indicates the runtime of *sPLINK*, which is the sum of the computation time in cohorts, network time to exchange the model parameters, aggregation time in the server, and idle time. During idle time, the aggregation result is ready but the cohorts are not aware of that since they periodically (every 10 seconds) ping the server. *sPLINK* computes the association test results for chi-square, linear regression, and logistic regression in 22 *min*, 56 *min*, and 3.5 *h*, respectively. Computation/aggregation time contributes the most/least to the total running time as expected (ignoring idle time). Compared to *Kamm et al.*^17^, *sPLINK* is 5 times faster for chi-square test (22 *min* vs. 110 *min*^2^) with less powerful hardware, larger sample size (5343 vs. 1080), and more number of SNPs (∼ 600*K* vs. ∼ 263*K*). In contrast to *Cho et al.*^18^, *sPLINK* is faster by a factor of 3 (3.5 *h* vs. ∼ 10.3 *h*^3^) comparing logistic regression test from *sPLINK* to Cochran–Armitage test implemented by that paper. Notice that logistic regression is a more complex algorithm than the Cochran–Armitage test for trend^43^.

Figure 9b depicts the network usage consumption of *sPLINK* for each association test. The cohorts and the server exchange total of 0.46 GB, 0.9 GB, and 8.33 GB traffic in chi-square, linear regression, and logistic regression, respectively. Logistic regression has higher volume of traffic exchange since the computation of beta coefficients are performed in an iterative fashion (maximum iteration count of 20). A fair comparison between *sPLINK* and SMPC-based frameworks from the network communication aspect is tricky. This is because network usage of *sPLINK* is independent of the sample size but linearly increases with the number of cohorts and SNPs while it only depends on the sample size and the number of SNPs in SMPC-based frameworks. However, in general, federated learning based approaches consume more network bandwidth than SMPC-based ones.

## Footnotes

1 *https://exbio.wzw.tum.de/splink*

2 The best result from *Kamm et al.*^17^ has been considered.

3 This value was computed based on the authors’ claim that their runtime linearly depends on the sample size and it takes 80 days to compute the results for a dataset with 1*M* individuals and 500*k* SNPs^18^.

## Notes

https://exbio.wzw.tum.de/splink

### Competing Interest Statement

The authors have declared no competing interest.

### Summary of Updates

Detailed comparison between sPLINK and meta-analysis for heterogeneous confounding factor scenario added; Runtime and network bandwidth usage results for sPLINK added; Figure 7 and Figure 8 revised; Author list updated; Concise comparison between sPLINK and state-of-the-art approaches including PLINK, meta-analysis, homomorphic encryption based methods and secure multiparty computing based frameworks added.

## References

1. Fareed, M. & Afzal, M. Single nucleotide polymorphism in genome-wide association of human population: A tool for broad spectrum service. Egypt. J. Med. Hum. Genet. 14, 123–134, https://doi.org/10.1016/j.ejmhg.2012.08.001 (2013).

2. Visscher, P. M. et al. 10 years of gwas discovery: biology, function, and translation. The Am. J. Hum. Genet. 101, 5–22 (2017).

3. Visscher, P. M., Brown, M. A., McCarthy, M. I. & Yang, J. Five years of gwas discovery. The Am. J. Hum. Genet. 90, 7–24 (2012).

4. De, R., Bush, W. S. & Moore, J. H. Bioinformatics challenges in genome-wide association studies (gwas). In Clinical Bioinformatics, 63–81 (Springer, 2014).

5. Purcell, S. et al. Plink: a tool set for whole-genome association and population-based linkage analyses. The Am. journal human genetics 81, 559–575 (2007).

6. Evangelou, E. & Ioannidis, J. P. Meta-analysis methods for genome-wide association studies and beyond. Nat. Rev. Genet. 14, 379–389 (2013).

7. Willer, C. J., Li, Y. & Abecasis, G. R. Metal: fast and efficient meta-analysis of genomewide association scans. Bioinformatics 26, 2190–2191 (2010).

8. Mägi, R. & Morris, A. P. Gwama: software for genome-wide association meta-analysis. BMC bioinformatics 11, 288 (2010).

9. Lunetta, K. L. Methods for meta-analysis of genetic data. Curr. protocols human genetics 77, 1–24 (2013).

10. Cantor, R. M., Lange, K. & Sinsheimer, J. S. Prioritizing gwas results: a review of statistical methods and recommendations for their application. The Am. J. Hum. Genet. 86, 6–22 (2010).

11. de Vlaming, R. et al. Meta-gwas accuracy and power (metagap) calculator shows that hiding heritability is partially due to imperfect genetic correlations across studies. PLoS genetics 13 (2017).

12. Gentry, C. Fully homomorphic encryption using ideal lattices. In Proceedings of the forty-first annual ACM symposium on Theory of computing, 169–178 (2009).

13. Cramer, R., Damgård, I. B. & Nielsen, J. B. Secure multiparty computation (Cambridge University Press, 2015).

14. McMahan, H. B., Moore, E., Ramage, D., Hampson, S. et al. Communication-efficient learning of deep networks from decentralized data. arXiv preprint arXiv:1602.05629 (2016).

15. Konečný, J. et al. Federated learning: Strategies for improving communication efficiency. arXiv preprint 1610.05492 (2016).

16. Shamir, A. How to share a secret. Commun. ACM 22, 612–613 (1979).

17. Kamm, L., Bogdanov, D., Laur, S. & Vilo, J. A new way to protect privacy in large-scale genome-wide association studies. Bioinformatics 29, 886–893 (2013).

18. Cho, H., Wu, D. J. & Berger, B. Secure genome-wide association analysis using multiparty computation. Nat. biotechnology 36, 547–551 (2018).

19. Shi, H. et al. Secure multi-party computation grid logistic regression (smac-glore). BMC medical informatics decision making 16, 89 (2016).

20. Alexandru, A. B. & Pappas, G. J. Secure multi-party computation for cloud-based control. Priv. Dyn. Syst. 179.

21. Lu, W.-J., Yamada, Y. & Sakuma, J. Privacy-preserving genome-wide association studies on cloud environment using fully homomorphic encryption. In BMC medical informatics and decision making, vol. 15, S1 (Springer, 2015).

22. Morshed, T., Alhadidi, D. & Mohammed, N. Parallel linear regression on encrypted data. In 2018 16th Annual Conference on Privacy, Security and Trust (PST), 1–5 (IEEE, 2018).

23. Kim, M., Song, Y., Wang, S., Xia, Y. & Jiang, X. Secure logistic regression based on homomorphic encryption: Design and evaluation. JMIR medical informatics 6, e19 (2018).

24. Chialva, D. & Dooms, A. Conditionals in homomorphic encryption and machine learning applications. arXiv preprint 1810.12380 (2018).

25. Wu, Y., Jiang, X., Kim, J. & Ohno-Machado, L. Grid binary logistic regression (glore): building shared models without sharing data. J. Am. Med. Informatics Assoc. 19, 758–764 (2012).

26. Jiang, W. et al. Webglore: a web service for grid logistic regression. Bioinformatics 29, 3238–3240 (2013).

27. Wang, S. et al. Expectation propagation logistic regression (explorer): distributed privacy-preserving online model learning. J. biomedical informatics 46, 480–496 (2013).

28. Friedman, J., Hastie, T. & Tibshirani, R. The elements of statistical learning, vol. 1 (Springer series in statistics New York, 2001).

29. Friedman, J., Hastie, T. & Tibshirani, R. The elements of statistical learning, vol. 1 (Springer series in statistics New York, 2001).

30. McHugh, M. L. The chi-square test of independence. Biochem. medica: Biochem. medica 23, 143–149 (2013).

31. Yang, Q., Liu, Y., Chen, T. & Tong, Y. Federated machine learning: Concept and applications. ACM Transactions on Intell. Syst. Technol. (TIST) 10, 1–19 (2019).

32. Kairouz, P. et al. Advances and open problems in federated learning. arXiv preprint 1912.04977 (2019).

33. Völzke, H. et al. Cohort profile: the study of health in pomerania. Int. journal epidemiology 40, 294–307 (2011).

34. Weiss, F. U. et al. Fucosyltransferase 2 (fut2) non-secretor status and blood group b are associated with elevated serum lipase activity in asymptomatic subjects, and an increased risk for chronic pancreatitis: a genetic association study. Gut 64, 646–656 (2015).

35. COPDGene. http://www.copdgene.org/ ((accessed Mar 22, 2020)).

36. Pillai, S. G. et al. A genome-wide association study in chronic obstructive pulmonary disease (copd): identification of two major susceptibility loci. PLoS genetics 5 (2009).

37. Pei, Y.-F., Tian, Q., Zhang, L. & Deng, H.-W. Exploring the major sources and extent of heterogeneity in a genome-wide association meta-analysis. Annals human genetics 80, 113–122 (2016).

38. EPACTS. https://genome.sph.umich.edu/wiki/EPACTS ((accessed Mar 22, 2020)).

39. Nguyen, T.-T., Huang, J. Z., Wu, Q., Nguyen, T. T. & Li, M. J. Genome-wide association data classification and snps selection using two-stage quality-based random forests. In BMC genomics, vol. 16, S5 (Springer, 2015).

40. Bellot, P., de los Campos, G. & Pérez-Enciso, M. Can deep learning improve genomic prediction of complex human traits? Genetics 210, 809–819 (2018).

41. Li, Y. R. & Keating, B. J. Trans-ethnic genome-wide association studies: advantages and challenges of mapping in diverse populations. Genome medicine 6, 91 (2014).

42. PLINK data formats. http://zzz.bwh.harvard.edu/plink/data.shtml ((accessed Mar 22, 2020)).

43. Donaldson, P., Daly, A., Ermini, L. & Bevitt, D. Genetics of complex disease (Garland Science, 2015).

